# Local adaptation contributes to gene expression divergence in maize

**DOI:** 10.1101/2020.08.01.231217

**Authors:** Jennifer Blanc, Karl A. G. Kremling, Edward Buckler, Emily B. Josephs

## Abstract

Gene expression links genotypes to phenotypes, so identifying genes whose expression is shaped by selection will be important for understanding the traits and processes underlying local adaptation. However, detecting local adaptation for gene expression will require distinguishing between divergence due to selection and divergence due to genetic drift. Here, we adapt a *Q_ST_* –*F_ST_* framework to detect local adaptation for transcriptome-wide gene expression levels in a population of diverse maize genotypes. We compare the number and types of selected genes across a wide range of maize populations and tissues, as well as selection on cold-response genes, drought-response genes, and coexpression clusters. We identify a number of genes whose expression levels are consistent with local adaptation and show that genes involved in stress-response show enrichment for selection. Due to its history of intense selective breeding and domestication, maize evolution has long been of interest to researchers, and our study provides insight into the genes and processes important for in local adaptation of maize.

## Introduction

Local adaptation occurs when different optimal trait values across environments lead to phenotypic differentiation among populations (Kawecki & Ebert, 2004). Identifying locally adapted traits is important for animal and crop production (Howden *et al.*, 2007; Takeda & Matsuoka, 2008), predicting response to climate change (Aitken *et al.*, 2008; Bay *et al.*, 2017; Franks & Hoffmann, 2012), and conservation genetics (Funk *et al.*, 2012). One commonly-used approach to identify local adaptation is *Q_ST_* –*F_ST_*, which tests for trait divergence (*Q_ST_*) that exceeds neutral expectations based on sequence divergence (*F_ST_*) (Spitze, 1993; Prout & Barker, 1993; Whitlock, 2008). However, while previous work has used *Q_ST_* –*F_ST_* and related approaches to identify specific traits showing evidence of selection, we lack broad-scale systematic investigations into the number and types of traits that are locally adapted.

Gene expression is a useful model trait for systematically investigating the evolutionary forces shaping phenotypic variation: expression is quantitative, can be heritable, and variation in gene expression can contribute to phenotypic variation and adaptation (Gibson & Weir, 2005; Roelofs *et al.*, 2006; Gilad *et al.*, 2006; Oleksiak *et al.*, 2002; Whitehead & Crawford, 2006; Gibson & Weir, 2005; Rockman & Kruglyak, 2006; Groen *et al.*, 2020). *Q_ST_* –*F_ST_* has previously identified local adaptation for gene expression in *D. melanogaster* and salmon (Roberge *et al.*, 2007; Kohn *et al.*, 2008) and a study has identified genes that showed relatively high or low *Q_ST_* in *Populus tremula* (Mähler *et al.*, 2017). Other studies have used an extension of *Q_ST_* –*F_ST_* developed by Ovaskainen *et al.* (2011) to identify genes showing evidence of local adaptation in expression (Leder *et al.*, 2015; Ravindran *et al.*, 2019). In this study, we leverage next generation sequencing data for expression and genetic variation to test for selection on expression of the entire transcriptome. In addition, we take advantage of a recent extension of *Q_ST_* –*F_ST_* that detects adaptation of continuous traits in large diversity panels that do not have clear subpopulations (Josephs *et al.*, 2019).

In this study, we investigate the role of local adaptation in shaping gene expression in the crop species *Zea mays.* Selection on gene expression has previously been shown to be important for maize evolution. For example, expression of the locus *tb1* (Doebley *et al.*, 1997; Wang *et al.*, 1999) is responsible for the evolution of apical dominance during domestication. Expression divergence is also prevalent between domesticated maize and its wild relative teosinte (Lemmon *et al.*, 2014) and expression variation in domesticated maize is often associated with phenotype (Kremling *et al.*, 2019). However deleterious mutations are important contributors to expression variation in maize (Kremling *et al.*, 2018), implying that not all expression variation in maize is adaptive.

Here we aim to understand the extent to which variation in gene expression in domesticated maize is driven by divergent selection caused by local adaptation and identify which genes show evidence of selection on their expression levels. We tested for selection using a published data set of 302 diverse maize lines each with RNAseq data from approximately 37,000 genes. We investigated enrichments of selective signals in genes that were differentially expressed in response to cold stress and drought, and selection on gene expression modules identified with coexpression network analyses taken from tissue-specific expression data. We detected selection on the expression of 60 unique genes across seven different tissue types and found an enrichment of drought-response genes among genes with the strongest signal of selection. Overall, these results show that local adaptation has shaped the expression of some genes and that this method has potential to identify specific genes and processes that are important for local adaptation.

## Methods

### Testing for selection on gene expression

Divergence between populations for a quantitative trait can be predicted by divergence at neutral genetic markers and additive genetic variation (*V_A_*), assuming the trait evolves neutrally and the trait value is made up of an additive combination of allelic effects (Henderson, 1950, 1953; Thompson, 2008). If a sample does not have discrete populations, the genetic principal components (PCs) that explain most of the genetic variation can be used as a measure of divergence between populations and the other PCs can be used to estimate *V_A_.* We briefly explain a test for selection using gene expression divergence measured across genetic PCs. More details on the test (*Q_PC_*) are available in Josephs *et al.* (2019).

Gene expression for a specific gene in *M* individuals is described by 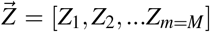. If the gene expression levels described by 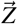 evolve neutrally, we can describe the distribution of 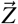 as follows:

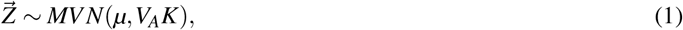

where *μ* is the mean expression value across individuals, *V_A_* is the additive genetic variation for expression, and *K* is the kinship matrix of the individuals. The kinship matrix *K* can be decomposed so that, *K* = *U*Λ*U^T^* where *U* is an *n x n* matrix where the columns are eigenvectors of *K* and Λ is a diagonal matrix of corresponding eigenvalues. The eigenvectors of *K* are the genetic principal components (PCs) of the population. We define 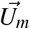 as the *m^th^* eigenvector and λ_m_ as the *m^th^* eigenvalue. The amount of trait variation explained by the *m^th^* PC, standardized by how much neutral genetic variation is explained by that PC, is

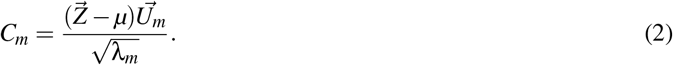

Under neutrality, *C_m_* ~ *N*(0, V*_A_*). If selection contributes to trait divergence along the *m^th^* PC, *C_m_* may fall outside the neutral distribution. For this study we tested the first five PCs for selection and the remaining PCs were used to estimate *V_A_*. To test for selection, we use a test statistic (*Q_PC_*).

For a focal PC *i*,

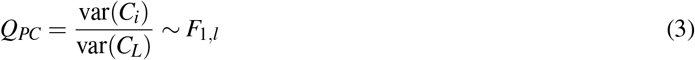

Intuitively, these ratios of variances are similar to a standard measure of *Q_ST_* in that the numerator describes between population expression level variance and the denominator describes within population expression level variance. Genes with a high value of *Q_PC_* will have expression levels are the most divergent at the between population level compared to the neutral expectation.

### Maize genomic and transcriptomic data

Expression and genotype data came from from a subset of a maize diversity panel generated by Flint-Garcia *et al.* (2005). These lines represent the diversity present in public-sector maize-breeding programs worldwide, including both temperate and tropical lines, as well as popcorn and sweet corn lines. Whole genome sequence (Bukowski *et al.*, 2017) and RNAseq data for 7 tissues (Kremling *et al.*, 2018) from plants grown in a common garden are available for these lines. Subsequent analysis only included genes that were expressed in all individuals for a given tissue type; which meant that we had between 8,435 and 11,555 genes per tissue type (Sample sizes listed in Table S1).

We used 78,342 randomly chosen SNPs to create a kinship matrix for each tissue type, reflecting the slightly differing set of lines present for each tissue. We arranged and standardized each kinship matrix so that each cell, *K_ij_* of the n x n matrix is the genotypic covariance between the *i^th^* and *j^ith^* lines following the procedure described in Josephs *et al.* (2019). After testing for selection as described above, FDR adjusted p-values were calculated to correct for multiple testing with the p.adjust function in R (Benjamini & Hochberg, 1995; R Core Team, 2020).

### Cluster Enrichment

We tested for local adaptation in the expression of gene coexpression modules. Walley *et al.* (2016) used weighted gene coexpression network analysis (WGCNA) to group genes that were similarly expressed in at least 4 tissues in one maize inbred line. This approach allowed them to group 31,447 mRNAs, 13,175 proteins, and 4,267 phosphoproteins into coexpression modules (clusters) and assign each cluster to the tissue(s) in which the cluster eigengene was most highly expressed. Their analysis resulted in 66 co-expression networks containing anywhere from 4 to 9574 genes. We calculated the median expression value for the genes in the 51 clusters that had more than 100 genes and used the same method outlined above on the median expression of each cluster to identify clusters that could be locally adapted.

### Environmental response genes

We tested for enrichment of signals of selection in genes that show expression changes in response to cold and drought. Cold-response genes were identified by Avila *et al.* (2018), who estimated the transcript abundance in leaves of 22,000 genes in two *Zea mays* inbred lines (CG60 and CG102) during and after cold temperature exposure and identified 10,549 genes differentially expressed in response to cold exposure. Drought-response genes were identified by Forestan *et al.* (2020), who measured transcript abundance in young leaves of the inbred line B73 and calculated differential expression between well-watered and drought stressed (10 days) treatments. Forestan *et al.* (2020) identified 3,181 differentially expressed genes (FDR < 0.01) and 28,983 non-differentially expressed genes.

Drought-response genes had higher daytime expression level in leaves than genes that didn’t show drought response (Figure S1). To ensure that overlaps between drought response genes and selected genes were not due to both sets of genes being biased towards high expression genes, we chose a subsample of 3500 of the non drought response genes with high expression to use as a comparison set (Figure S1). There was not a significant difference in daytime leaf expression level between cold response and non cold response genes, so we did not adjust the test for gene expression level.

With both datasets, we used a Fisher’s exact test to compare the proportion of genes that show evidence of selection (un-adjusted p value less than 0.05) in environmental-response genes compared with other genes (see Tables S2, S3, and S4 for sample sizes). We used the un-adjusted p value so that we had enough genes in each category to use Fisher’s exact test. We only tested for enrichment in tissue-PC combinations that had evidence of at least one selected gene at FDR < 0.1. P values were then adjusted for multiple testing using a Bonferroni correction (n=15).

### GO Enrichment Analysis

We tested subsets of genes identified as having signals of selection on gene expression for enrichment of GO biological process terms using the GO Enrichment Analysis tool on geneontology.org. (Ashburner *et al.*, 2000; Consortium, 2019; Mi *et al.*, 2019) We used the genes that went into our selection analysis for a given tissue as the reference list and the genes whose expression was under selection along a specific PC in that same tissue as the analyzed list. We used Fisher’s exact test and FDR as calculated by the Benjamini-Hochberg procedure for multiple testing correction as the settings for the enrichment analysis.

## Results

### Detecting selection on expression of individual genes

We tested for selection on gene expression of 8,435 to 11,555 genes in seven tissues for 109 to 239 genotypes (see Table S1 for sample sizes), along the first five PCs within each tissue type. Note that because there were different genotypes sampled in each tissue type, the genetic PCs do not always correspond across tissues (Figures S3, S4, S5). Across all tissues, PC 1 separated out tropical from temperate genotypes and lower PCs separated stiff stalk from non stiff stalk genotypes, popcorns from other genotypes, or separated out genotypes within the stiff stalk and/or non stiff stalk subpopulations (Figures S3, S4, S5).

Sixty unique genes show evidence of expression divergence consistent with local adaptation along one of the first 5 PCs (FDR < 0.1, Figure 1A). We plot an example of the signal of selection on two genes to demonstrate what expression values look like when selection is inferred along a specific PC (Figure 1B,C). There were 5 genes that had evidence for selection on expression in multiple tissues and/or multiple PCs. The PC-tissue combination with the most genes under selection was PC 5 in adult leaf expression measured during the day. Genes with divergence along PC 5 in adult leaf tissue are enriched for GO biological process terms cellulose catabolic process (FDR = 0.0323), plant-type cell wall biogenesis (FDR = 0.00853), and glucan biosynthetic process (FDR = 0.0287).

**Figure 1:**
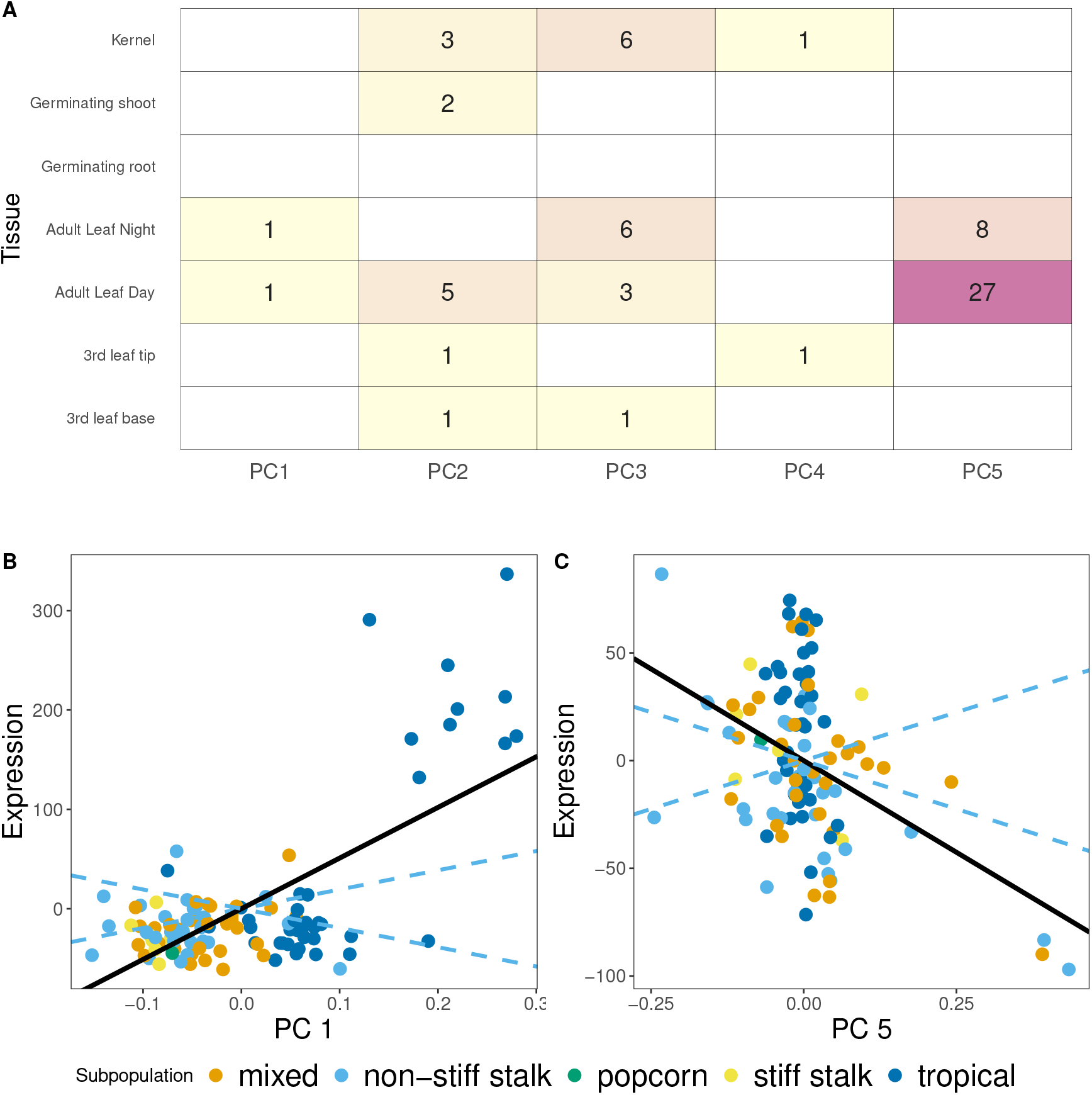
Signals of selection on gene expression in domesticated maize. A) The number of genes where *FDR* < 0.1 in each of the 7 tissues for the first 5 PCs. B) PC 1 plotted against the mean-centered expression level of the gene GRMZM2G152686 as expressed in adult leaves during the day. Each point represents one maize genotype and is colored by subpopulation. The solid line shows the linear regression and the dashed lines show 95% confidence intervals of the neutral expectation. C) Similar to plot (B) except PC 5 plotted against mean centered expression of the gene GRMZM2G069762

### Selection on expression of coexpression clusters

Gene expression is often correlated across genes, so summarizing expression across coexpression clusters could improve power to detect selection (Kliebenstein, 2020). With this in mind, we summarized expression across previously identified coexpression modules (Walley *et al.*, 2016) and tested for selection on median gene expression for each module. However, none of the clusters showed evidence of selection (*FDR >* 0.1). The test with the strongest evidence of selection was the ‘Root Meristem’ cluster, which showed evidence of selection along PC 5 in leaf adult tissue measured during the day (*p* = 2.4×10^−4^, FDR = 0.43). While the ‘Root Meristem’ cluster had the highest expression in root meristems in Walley *et al.* (2016), many of these genes were still expressed in adult leaves in their study. Overall, these results suggest that coexpression clusters, as identified by correlations in expression within one genotype, are not broad targets of selection.

### Selection on expression of environmental response genes

The spread of maize into North America required adaptation to different climatic factors (Swarts *et al.*, 2017), so we investigated selection specifically on genes that were differentially expressed in response to cold (Avila *et al.*, 2018) and in response to drought (Forestan *et al.*, 2020).

To test for evidence of selection on genes that were differentially expressed in response to cold, we compared selection signals in 12,239 genes that showed differential expression (*FDR* < 0.1) after either one or four days of cold treatment to 11,379 genes that did not show evidence of differential expression using data from Avila *et al.* (2018). We only investigated the 15 tissue-PC combinations where at least one gene showed significant evidence of selection at FDR < 0.01. The strongest signal for enrichment was for daytime expression in adult leaf tissue along PC 5, where genes whose expression changed in response to cold were more likely to have evidence of local adaptation for expression (p Bonferonni p = 0.06, Table S2, Figure S2).

We found a significant enrichment of selection signals in 560 genes that showed decreased expression in response to drought in the B73 line compared to 3,500 genes with similar leaf expression levels but that were not differentially expressed in drought (Table S3). Specifically, expression in adult leaf tissue in both day and night showed evidence of enrichment for signals of selection along PC 5. 14% of genes down-regulated in drought showed evidence of selection on leaf expression during day and night, while 8.1% of genes without drought response had evidence of selection for leaf expression during the day and 6.9% had evidence for selection on leaf expression at night (Bonferroni p = 0.00363 for day Bonferroni p = 1.635×10^−5^for night) (Figure 2). The 328 genes that had increased expression in drought did not show any enrichment for selection (Figure 2, Table S4).

**Figure 2:**
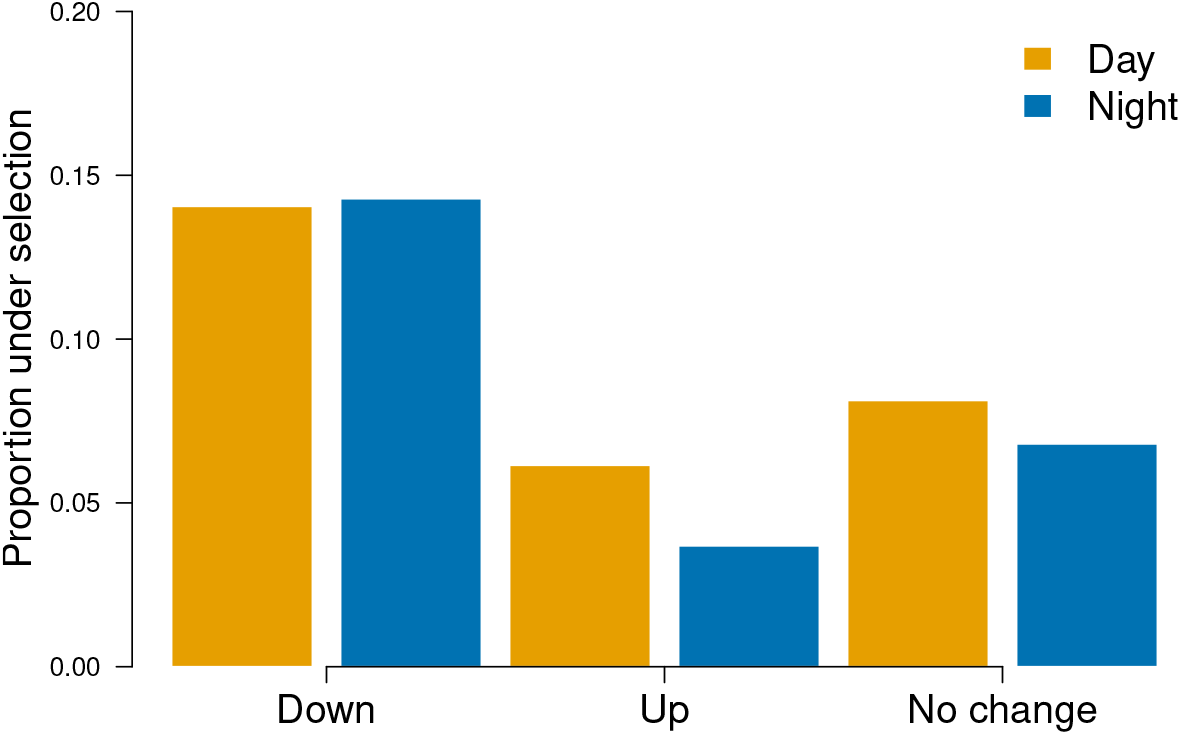
(A) Enrichment for signals of selection in genes down-regulated in drought. The percentage of genes that show evidence of selection along PC 5 (*p* < 0.05) in adult leaf expression during the day (orange) and night (blue) for genes that are down-regulated in drought, up-regulated in drought, and show no change in response to drought.

## Discussion

Systematically identifying genes important for local adaptation is crucial for understanding how local adaptation shapes trait variation. Here, we used an extension of *Q_ST_* –*F_ST_* to identify genes with expression divergence consistent with local adaptation in domesticated maize. Out of a dataset of expression of ~ 10,000 genes measured across seven tissue types, we identified 60 genes with expression divergence consistent with local adaptation in at least one tissue type. Additionally, we found evidence that genes involved in drought response and cold response are enriched for signals of selection.

Our results contribute to a growing body of evidence that genetic variation for gene expression is shaped by selection. Previous studies in maize and other species have shown that rare variants affecting gene expression are often under negative selection (Kremling *et al.*, 2018; Josephs *et al.*, 2015; Glassberg *et al.*, 2019) and that there is weak stabilizing selection on gene expression levels in the field (Groen *et al.*, 2020). Alongside evidence for negative selection, *Q_ST_* –*F_ST_* and related analyses have demonstrated that local adaptation shapes between-population divergence in expression for some genes (Whitehead & Crawford, 2006; Kohn *et al.*, 2008; Roberge *et al.*, 2007; Ravindran *et al.*, 2019; Jueterbock *et al.*, 2016). This is the first study to use *Q_PC_,* a *Q_ST_* –*F_ST_*-based method that detects selection on expression in the absence of clear subpopulations. With increasing availability of large transcriptomic studies conducted on diversity panels, methods for detecting selection on expression in the absence of clear subpopulations will be useful for understanding how selection shapes expression variation.

The enrichment of signals of adaptive divergence in genes involved in environmental response provides evidence for types of environmental factors that could contribute to adaptive divergence in expression. A number of pieces of evidence suggest that genes important for drought response had expression values shaped by local adaptation. There is an enrichment for signals of selection along PC 5 in genes that have decreased expression in response to experimental drought. One gene that shows adaptive expression divergence along PC 5 in leaf tissue (FDR = 0.02 for day and FDR = 0.01 for night) codes for the protein ZmRD22B, a putative maize RD22-like protein (Phillips & Ludidi, 2017). RD22 proteins are thought to play a role in drought response through the ABA (abscisic acid) signalling pathway (Xu *et al.*, 2010) and ZmRD22B itself is predicted to localize to the cell wall and is upregulated in response to drought and exogenous ABA (Phillips & Ludidi, 2017). Additionally, the group of genes we detected as having significant expression divergence along PC 5 in leaf tissue, including ZmRD22B, are enriched for GO biological processes cellulose catabolic process, plant-type cell wall biogenesis, and glucan biosynthetic process. In leaf tissue, PC 5 separated out individuals in the non-stiff-stalk heterotic group of maize, suggesting that further investigations into gene expression and drought response in this subpopulation may be a promising future direction.

However, the link between genes important for stress response and evidence of local adaptation for gene expression in well-watered conditions is complex. The environmental response genes used in this study were identified from studies of differential expression in a few temperate maize genotypes. Stress-induced changes in gene expression could be beneficial responses that help the individual cope with stress or deleterious responses caused by the individual’s inability to maintain function in stressful conditions (Ghalambor *et al.*, 2007). If stress responses tend to be adaptive and improve function in the stressful condition, then local adaptation for expression in non-stressful conditions could reflect constitutive changes in expression in genotypes more likely to experience the stress. In contrast, if stress responses tend to be maladaptive in the stress environment, then local adaptation for expression in non-stressed environment could reflect further selection for reduced response even in non-stressful environments. For both cases, clearly understanding selection on the expression of environment-response genes will require additional experiments that measure expression changes in different environments across a diverse panel of genotypes.

While our method was successful in identifying genes whose expression is consistent with local adaptation, we only detected selection on 60 genes. Maize domestication and improvement has involved genome-wide selection (Wright *et al.*, 2005; Hufford *et al.*, 2012; Wang *et al.*, 2020; Swarts *et al.*, 2017), so we may expect to see evidence of selection on the expression on many more than 60 genes. There are a few potential explanations for why evidence of selection on gene expression may be limited. First, transcriptomes are a snapshot in a specific developmental time and environment and this study may have missed tissues, developmental time points, or environments in which expression has been under strong selection. Second, *Q_PC_* loses power when there is high environmental variation (*V_E_*) for a trait. *V_E_* increases trait variance explained by later (’within population’) PCs and, since these later PCs are used to generate a neutral expectation of divergence along focal PCs, high *V_E_* will increase the amount of expression variation expected under neutrality (Figure S6). Overall, this means that high *V_E_* will reduce power to detect selection (Josephs *et al.*, 2019). This reduction in power due to *V_E_* may be especially strong in expression data, which tends to be noisy and measured in few or no replicates.

An additional limitation of this study and the *Q_PC_* approach is that we were only able to investigate genes that were expressed in all individuals for a given tissue type. *Q_PC_* models phenotypes as additive combinations of allelic effects (Josephs *et al.*, 2019), and so the model is not robust to phenotypic distributions where a large number of individuals have a phenotype of 0. However, many of the expression changes that are important for phenotypic change may involve genes being turned on and off, not quantitative expression changes (Zhou *et al.*, 2020). In addition, maize has many presence-absence variants and the expression of these genes will appear to be 0 in individuals with the absent allele (Zhou *et al.*, 2019; Hirsch *et al.*, 2014). Methods to detect adaptive divergence in traits with non-normal distributions will be useful for future progress and may be able to detect more instances of adaptation.

Altogether, our work demonstrates that *Q_PC_* can be used to systematically detect genes whose expression is shaped by local adaptation and has shown its effectiveness in a large dataset from domesticated maize. We not only were able to detect selection on specific genes, but on combinations of genes based on environmental response patterns. Overall, our work shows that this method has potential for use in a number of large diversity panels while suggesting ways forward for better detecting selection on gene expression.

## Acknowledgements

We thank Jeremy Berg, Graham Coop, Jeff Ross-Ibarra, Nathan Springer, and members of the Coop, Ross-Ibarra, Novembre, Berg, and Steinrücken labs for discussions and helpful feedback. This work was funded by NSF IOS-1523733 and NSF-IOS-1934384 to EBJ, and NSF-IOS-1546719 to Graham Coop and Jeffrey Ross-Ibarra.

---

**Figure S1:**
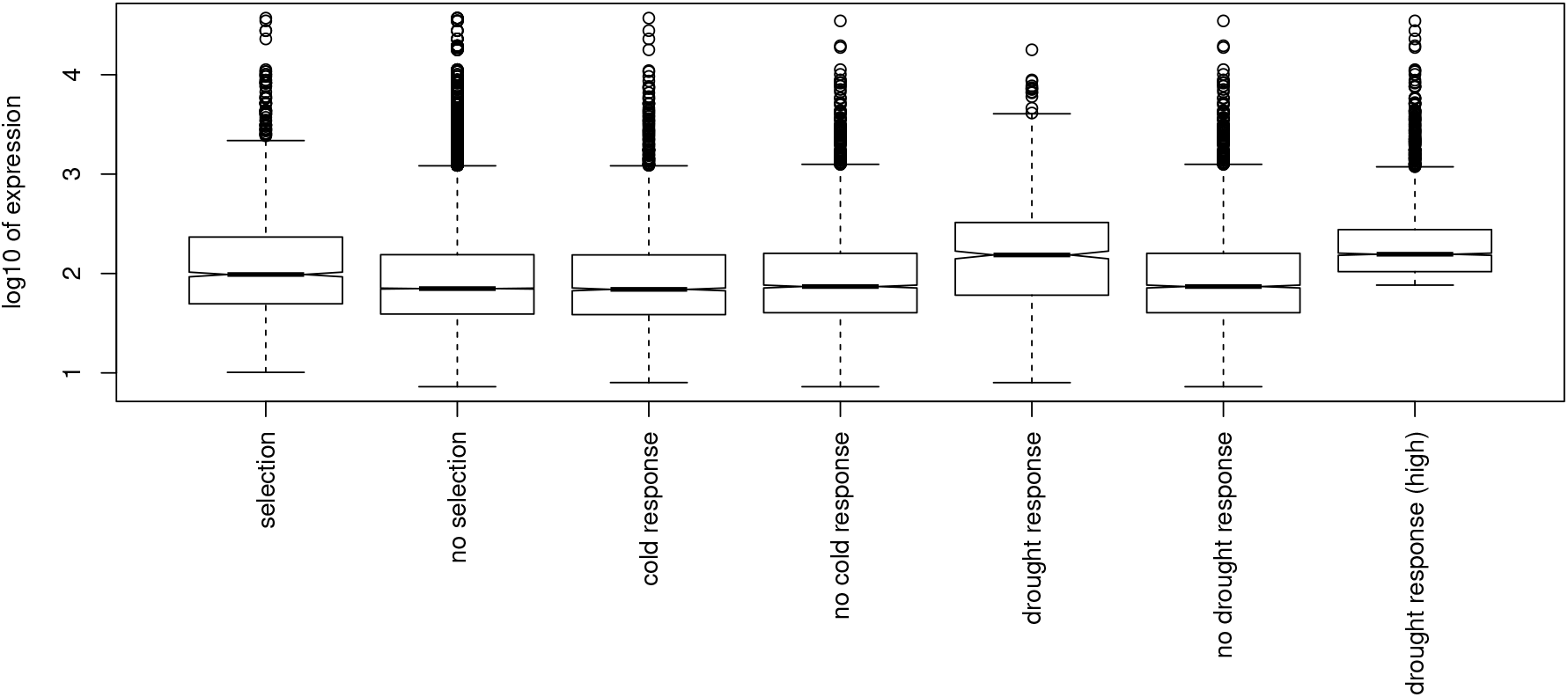
Expression level of genes in different categories

**Table S1:** Number of genes and individuals used to test for selection in each tissue

| Tissue | Gene Number | Individual Number |
| --- | --- | --- |
| Kernel | 9,814 | 207 |
| Germinating shoot | 10,195 | 239 |
| Germinating root | 10,500 | 232 |
| Adult leaf night | 8,435 | 110 |
| Adult leaf day | 8,879 | 109 |
| 3<sup>rd</sup> leaf tip | 8,489 | 237 |
| 3<sup>rd</sup> leaf base | 11,555 | 236 |

**Table S2:**
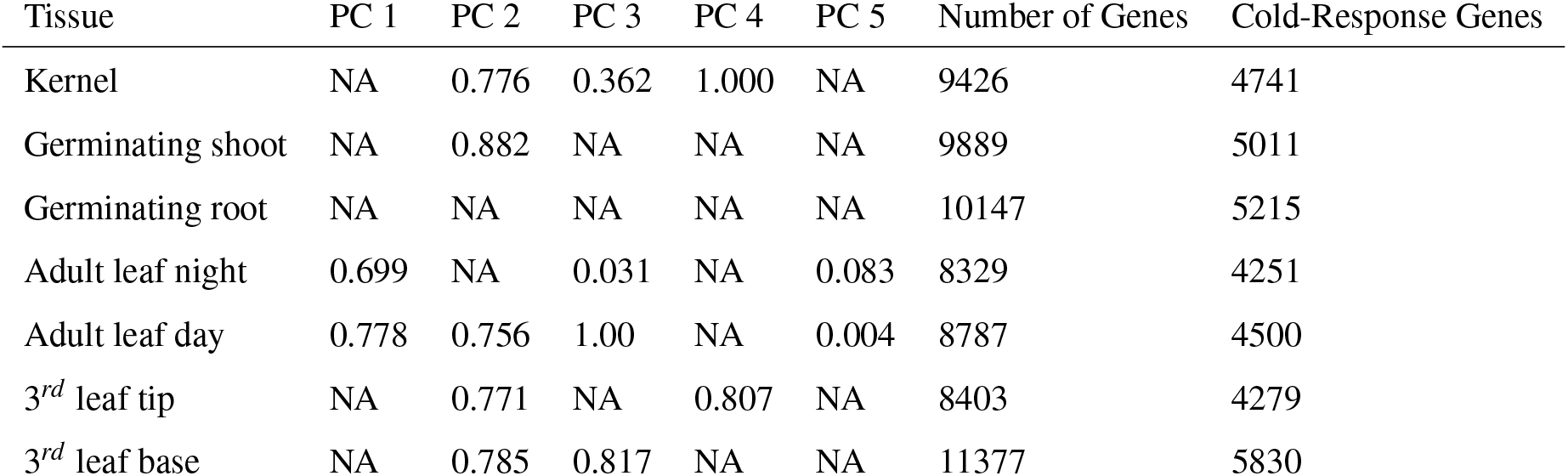
Uncorrected p-values and sample sizes for chi-squared test for enrichment of signals of selection in cold-response genes. P-values only shown for PC/tissue combinations with at least 1 significantly selected gene (FDR < 0.1).

| Tissue | PC 1 | PC 2 | PC 3 | PC 4 | PC 5 | Number of Genes | Cold-Response Genes |
| --- | --- | --- | --- | --- | --- | --- | --- |
| Kernel | NA | 0.776 | 0.362 | 1.000 | NA | 9426 | 4741 |
| Germinating shoot | NA | 0.882 | NA | NA | NA | 9889 | 5011 |
| Germinating root | NA | NA | NA | NA | NA | 10147 | 5215 |
| Adult leaf night | 0.699 | NA | 0.031 | NA | 0.083 | 8329 | 4251 |
| Adult leaf day | 0.778 | 0.756 | 1.00 | NA | 0.004 | 8787 | 4500 |
| 3<sup>rd</sup> leaf tip | NA | 0.771 | NA | 0.807 | NA | 8403 | 4279 |
| 3<sup>rd</sup> leaf base | NA | 0.785 | 0.817 | NA | NA | 11377 | 5830 |

**Figure S2:**
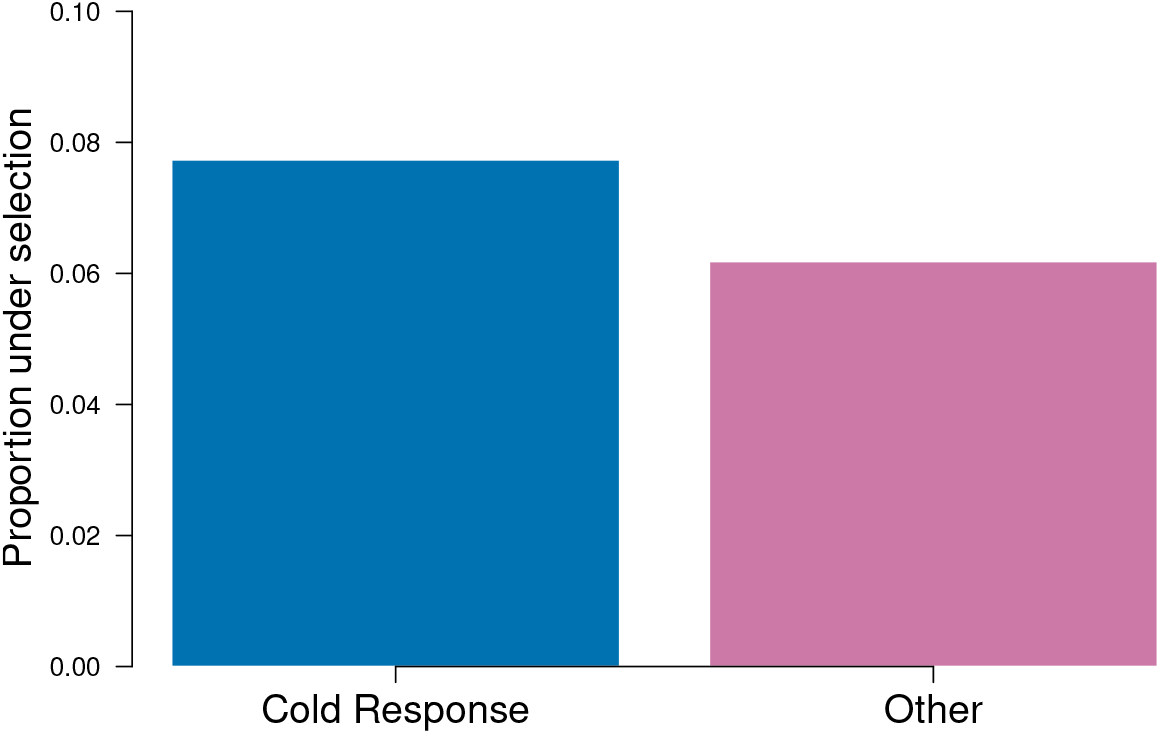
Enrichment for signals of selection in genes with differential response to cold treatment. The percentage of genes that show evidence of selection along PC 5 (*p* < 0.05) in adult leaf expression during the day for genes that have expression change in cold and no change in response to cold. While there is a slight enrichment of signals of selection in cold-response genes, this enrichment is not significant after a Bonferroni correction for multiple testing (*p* = 0.09)

**Figure S3:**
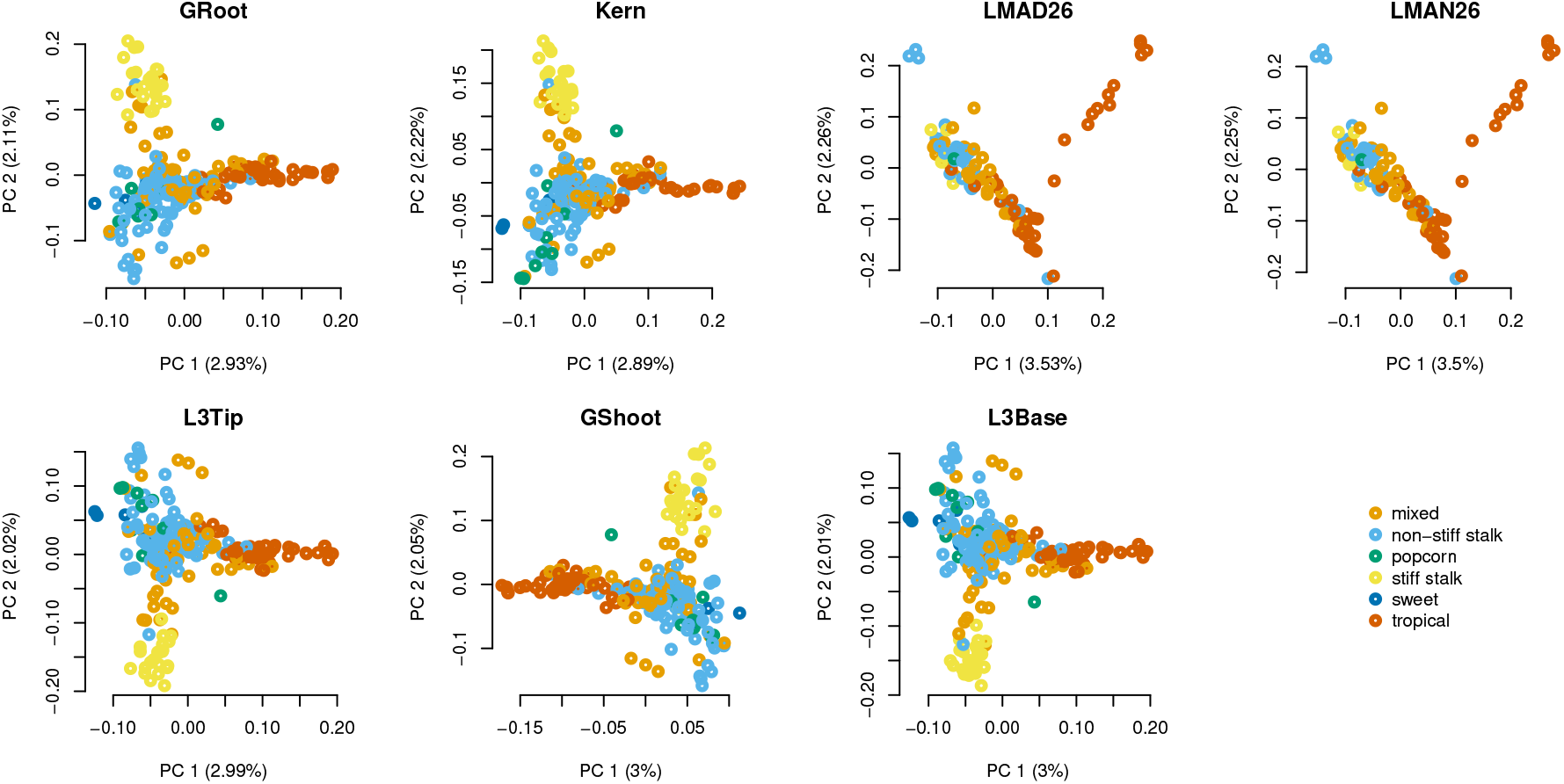
The first two genetic PCs of genotypes in each tissue expression dataset. Each point represents one genotype, colored by subpopulation. The x axis is PC 1 and the Y axis is PC 2, labeled by the percentage of variation that each PC explains.

**Figure S4:**
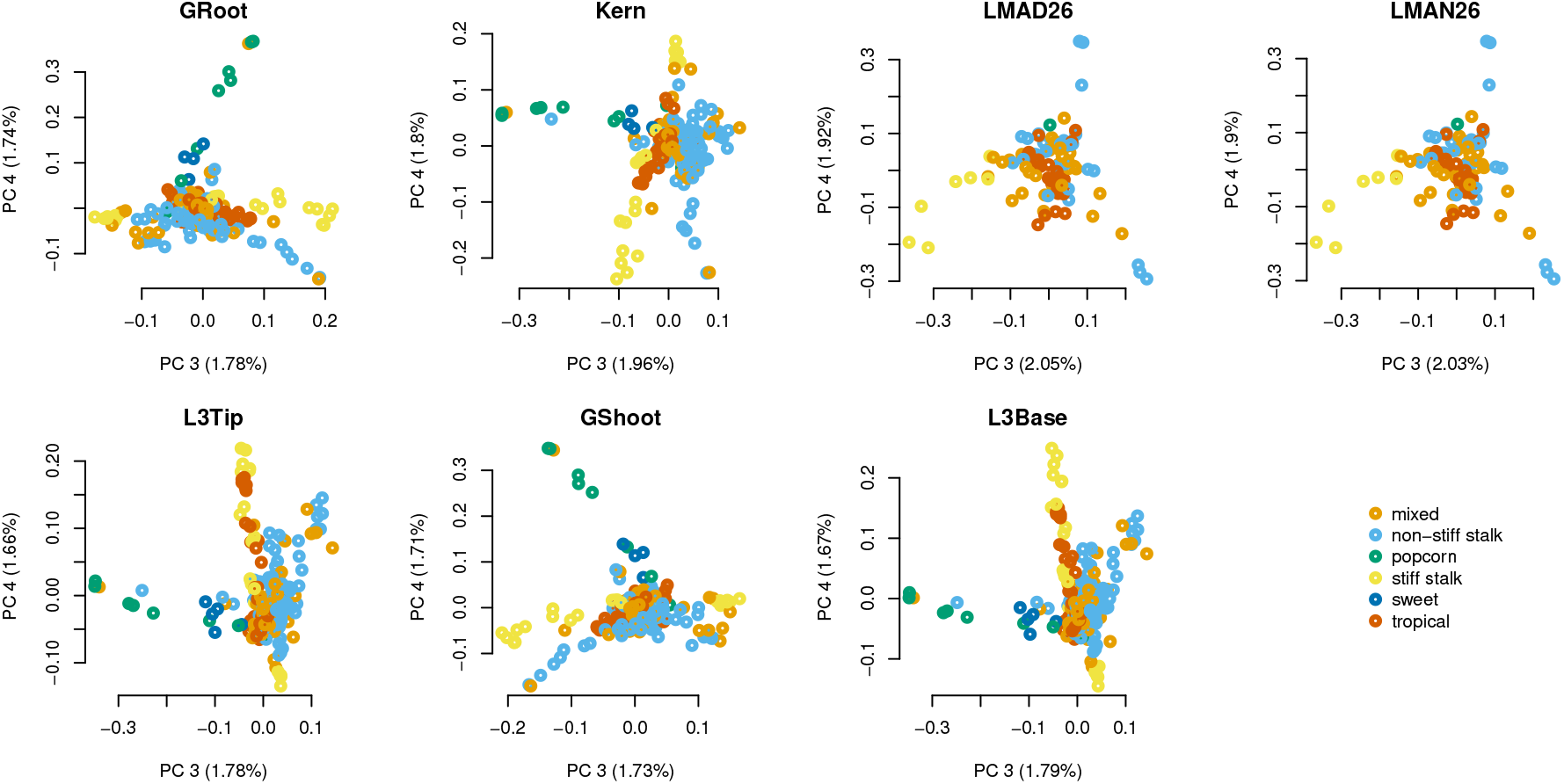
The third and fourth genetic PCs of genotypes in each tissue expression dataset. Each point represents one genotype, colored by subpopulation. The x axis is PC 3 and the Y axis is PC 4, labeled by the percentage of variation that each PC explains.

**Figure S5:**
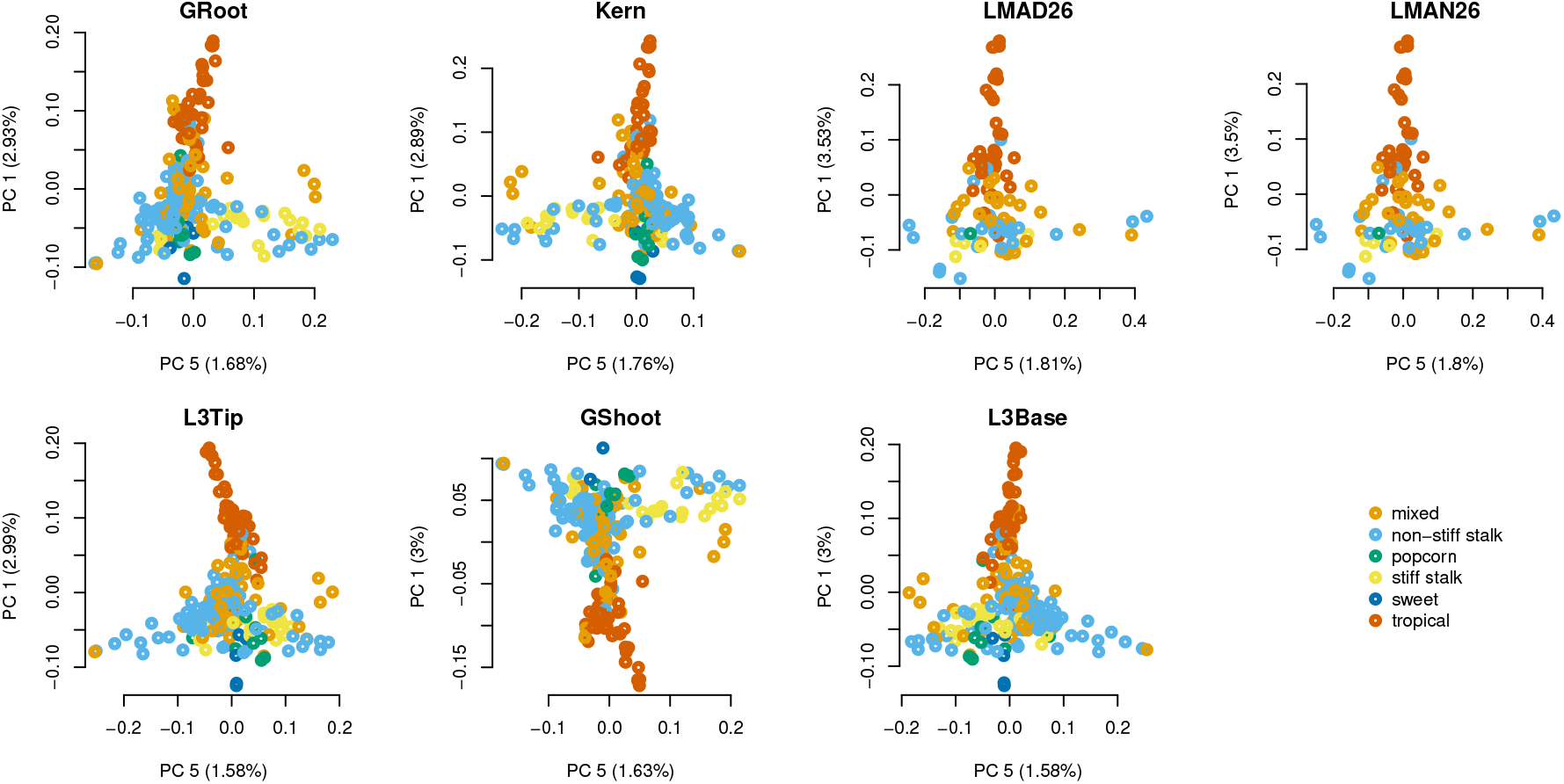
The first and fifth genetic PCs of genotypes in each tissue expression dataset. Each point represents one genotype, colored by subpopulation. The x axis is PC 5 and the Y axis is PC 1, labeled by the percentage of variation that each PC explains.

**Figure S6:**
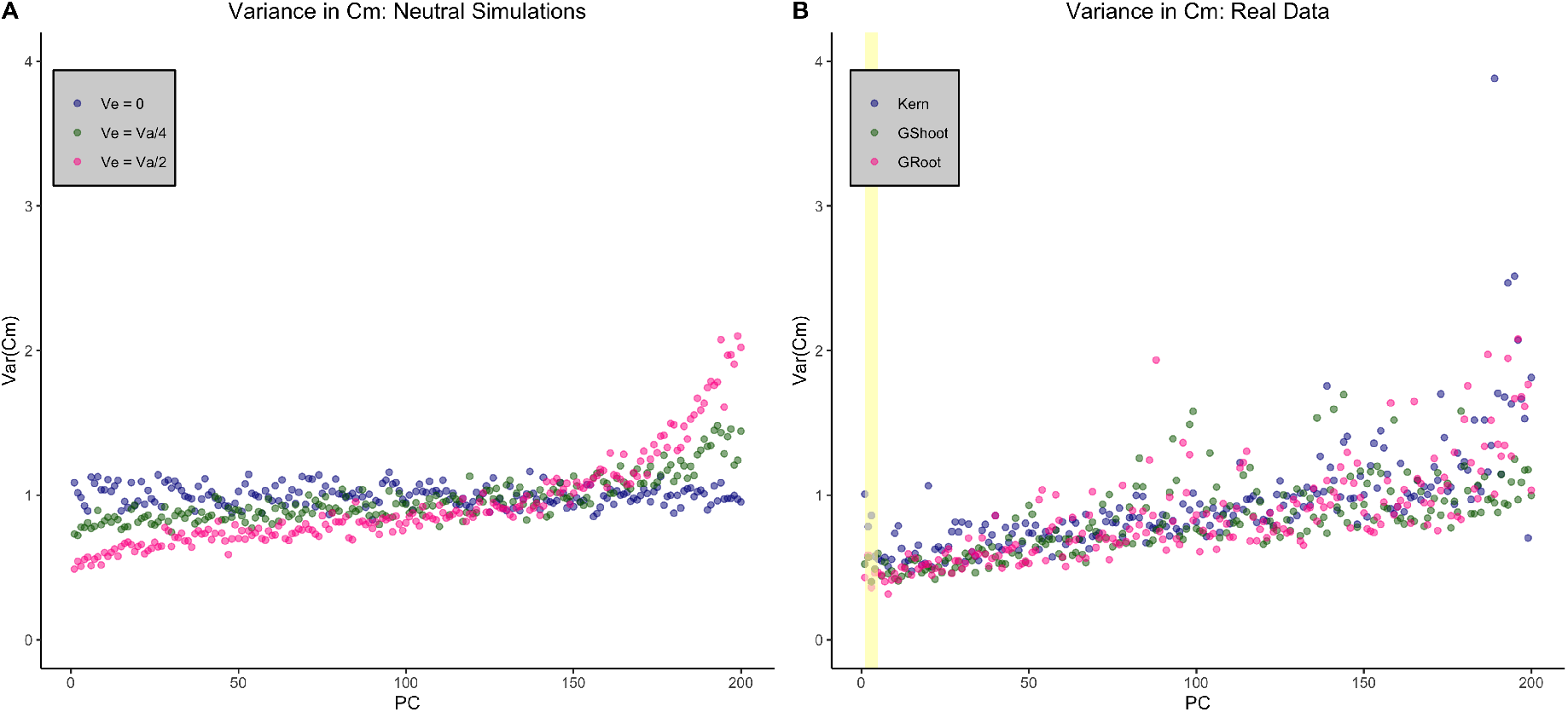
(A) Variance in *C_m_* values for neutral simulations with different levels of environmental variance using the kinship matrix generated from the 207 Kernel lines. (B) Variance in *C_m_* values for actual expression values for 3 different tissue types. The yellow box highlights the five 5 PCs along which expression divergence was tested.

**Table S3:** Uncorrected p-values for chi-squared test for enrichment of signals of selection in down-regulated drought-response genes. P-values only shown for PC/tissue combinations with at least 1 significantly selected gene (FDR < 0.1).

| Tissue | PC 1 | PC 2 | PC 3 | PC 4 | PC 5 | Number of Genes | Down-Regulated Drought-Response Gene |
| --- | --- | --- | --- | --- | --- | --- | --- |
| Kernel | NA | 0.391 | 0.878 | 0.023 | NA | 3031 | 352 |
| Germinating shoot | NA | 0.696 | NA | NA | NA | 3450 | 463 |
| Germinating root | NA | NA | NA | NA | NA | 3045 | 366 |
| Adult leaf night | 0.083 | NA | 0.344 | NA | 0.0000109 | 3605 | 464 |
| Adult leaf day | 0.198 | 0.033 | 0.639 | NA | 0.000242 | 4065 | 566 |
| 3<sup>rd</sup> leaf tip | NA | 0.720 | NA | 0.451 | NA | 3735 | 476 |
| 3<sup>rd</sup> leaf base | NA | 0.034 | 0.5723 | NA | NA | 3687 | 511 |

**Table S4:** Uncorrected p-values and sample sizes for chi-squared test for enrichment of signals of selection in up-regulated drought-response genes. P-values only shown for PC/tissue combinations with at least 1 significantly selected gene (FDR < 0.1).

| Tissue | PC 1 | PC 2 | PC 3 | PC 4 | PC 5 | Number of Genes | Up-Regulated Drought-Response Genes |
| --- | --- | --- | --- | --- | --- | --- | --- |
| Kernel | NA | 0.689 | 0.051 | 0.124 | NA | 2844 | 165 |
| Germinating shoot | NA | 0.181 | NA | NA | NA | 3142 | 155 |
| Germinating root | NA | NA | NA | NA | NA | 2679 | 161 |
| Adult leaf night | 0.379 | NA | 1.000 | NA | 0.077 | 3394 | 253 |
| Adult leaf day | 0.337 | 0.915 | 0.104 | NA | 0.319 | 3827 | 328 |
| 3<sup>rd</sup> leaf tip | NA | 0.434 | NA | 0.305 | NA | 3513 | 254 |
| 3<sup>rd</sup> leaf base | NA | 0.485 | 1.000 | NA | NA | 3337 | 161 |

## References

Aitken SN, Yeaman S, Holliday JA, Wang T, Curtis-McLane S. 2008. Adaptation, migration or extirpation: climate change outcomes for tree populations. Evolutionary applications, 1: 95–111.

Ashburner M, Ball CA, Blake JA, Botstein D, Butler H, Cherry JM, Davis AP, Dolinski K, Dwight SS, Eppig JT et al. 2000. Gene ontology: tool for the unification of biology. Nature genetics, 25: 25–29.

Avila LM, Obeidat W, Earl H, Niu X, Hargreaves W, Lukens L. 2018. Shared and genetically distinct zea mays transcriptome responses to ongoing and past low temperature exposure. BMC genomics, 19: 761.

Bay RA, Rose N, Barrett R, Bernatchez L, Ghalambor CK, Lasky JR, Brem RB, Palumbi SR, Ralph P. 2017. Predicting responses to contemporary environmental change using evolutionary response architectures. The American Naturalist, 189: 463–473.

Benjamini Y, Hochberg Y. 1995. Controlling the false discovery rate: a practical and powerful approach to multiple testing. Journal of the Royal statistical society: series B (Methodological), 57: 289–300.

Bukowski R, Guo X, Lu Y, Zou C, He B, Rong Z, Wang B, Xu D, Yang B, Xie C et al. 2017. Construction of the third-generation zea mays haplotype map. Gigascience, 7: gix134.

Consortium GO. 2019. The gene ontology resource: 20 years and still going strong. Nucleic acids research, 47: D330–D338.

Doebley J, Stec A, Hubbard L. 1997. The evolution of apical dominance in maize. Nature, 386: 485.

Flint-Garcia SA, Thuillet AC, Yu J, Pressoir G, Romero SM, Mitchell SE, Doebley J, Kresovich S, Goodman MM, Buckler ES. 2005. Maize association population: a high-resolution platform for quantitative trait locus dissection. The Plant Journal, 44: 1054–1064.

Forestan C, Farinati S, Zambelli F, Pavesi G, Rossi V, Varotto S. 2020. Epigenetic signatures of stress adaptation and flowering regulation in response to extended drought and recovery in zea mays. Plant, cell & environment, 43: 55–75.

Franks SJ, Hoffmann AA. 2012. Genetics of climate change adaptation. Annual review of genetics, 46.

Funk WC, McKay JK, Hohenlohe PA, Allendorf FW. 2012. Harnessing genomics for delineating conservation units. Trends in ecology & evolution, 27: 489–496.

Ghalambor CK, McKay JK, Carroll SP, Reznick DN. 2007. Adaptive versus non-adaptive phenotypic plasticity and the potential for contemporary adaptation in new environments. Functional ecology, 21: 394–407.

Gibson G, Weir B. 2005. The quantitative genetics of transcription. Trends in Genetics, 21: 616 – 623.

Gilad Y, Oshlack A, Smyth GK, Speed TP, White KP. 2006. Expression profiling in primates reveals a rapid evolution of human transcription factors. Nature, 440: 242.

Glassberg EC, Gao Z, Harpak A, Lan X, Pritchard JK. 2019. Evidence for weak selective constraint on human gene expression. Genetics, 211: 757–772.

Groen SC, Ćalić I, Joly-Lopez Z, Platts AE, Choi JY, Natividad M, Dorph K, Mauck WM, Bracken B, Cabral CLU et al. 2020. The strength and pattern of natural selection on gene expression in rice. Nature, 578: 572–576.

Henderson CR. 1950. Estimation of genetic parameters. In: Biometrics. INTERNATIONAL BIOMETRIC SOC 14411 ST, NW, SUITE 700, WASHINGTON, DC 20005-2210, vol. 6, pp. 186–187.

Henderson CR. 1953. Estimation of variance and covariance components. Biometrics, 9: 226–252.

Hirsch CN, Foerster JM, Johnson JM, Sekhon RS, Muttoni G, Vaillancourt B, Peñxagaricano F, Lindquist E, Pedraza MA, Barry K et al. 2014. Insights into the maize pan-genome and pan-transcriptome. The Plant cell, 26: 121–135.

Howden SM, Soussana JF, Tubiello FN, Chhetri N, Dunlop M, Meinke H. 2007. Adapting agriculture to climate change. Proceedings of the national academy of sciences, 104: 19691–19696.

Hufford MB, Xu X, Van Heerwaarden J, Pyhäjärvi T, Chia JM, Cartwright RA, Elshire RJ, Glaubitz JC, Guill KE, Kaeppler SM et al. 2012. Comparative population genomics of maize domestication and improvement. Nature genetics, 44: 808.

Josephs EB, Berg JJ, Ross-Ibarra J, Coop G. 2019. Detecting adaptive differentiation in structured populations with genomic data and common gardens. Genetics, 211: 989–1004.

Josephs EB, Lee YW, Stinchcombe JR, Wright SI. 2015. Association mapping reveals the role of purifying selection in the maintenance of genomic variation in gene expression. Proceedings of the National Academy of Sciences, 112: 15390–15395.

Jueterbock A, Franssen SU, Bergmann N, Gu J, Coyer JA, Reusch TB, Bornberg-Bauer E, Olsen JL. 2016. Phylogeographic differentiation versus transcriptomic adaptation to warm temperatures in zostera marina, a globally important seagrass. Molecular ecology, 25: 5396–5411.

Kawecki TJ, Ebert D. 2004. Conceptual issues in local adaptation. Ecology letters, 7: 1225–1241.

Kliebenstein DJ. 2020. Using networks to identify and interpret natural variation. Current Opinion in Plant Biology, 54: 122–126.

Kohn MH, Shapiro J, Wu CI. 2008. Decoupled differentiation of gene expression and coding sequence among drosophila populations. Genes & genetic systems, 83: 265–273.

Kremling KA, Chen SY, Su MH, Lepak NK, Romay MC, Swarts KL, Lu F, Lorant A, Bradbury PJ, Buckler ES. 2018. Dysregulation of expression correlates with rare-allele burden and fitness loss in maize. Nature, 555: 520.

Kremling KA, Diepenbrock CH, Gore MA, Buckler ES, Bandillo NB. 2019. Transcriptome-wide association supplements genome-wide association in zea mays. G3: Genes, Genomes, Genetics, pp. g3–400549.

Leder EH, McCairns RS, Leinonen T, Cano JM, Viitaniemi HM, Nikinmaa M, Primmer CR, Merilä J. 2015. The evolution and adaptive potential of transcriptional variation in sticklebackssignatures of selection and widespread heritability. Molecular biology and evolution, 32: 674–689.

Lemmon ZH, Bukowski R, Sun Q, Doebley JF. 2014. The role of cis regulatory evolution in maize domestication. PLoS genetics, 10: e1004745.

Mähler N, Wang J, Terebieniec BK, Ingvarsson PK, Street NR, Hvidsten TR. 2017. Gene co-expression network connectivity is an important determinant of selective constraint. PLoS genetics, 13: e1006402.

Mi H, Muruganujan A, Ebert D, Huang X, Thomas PD. 2019. Panther version 14: more genomes, a new panther go-slim and improvements in enrichment analysis tools. Nucleic acids research, 47: D419–D426.

Oleksiak MF, Churchill GA, Crawford DL. 2002. Variation in gene expression within and among natural populations. Nature genetics, 32: 261.

Ovaskainen O, Karhunen M, Zheng C, Arias JMC, Merila J. 2011. A new method to uncover signatures of divergent and stabilizing selection in quantitative traits. Genetics, 189: 621–632.

Phillips K, Ludidi N. 2017. Drought and exogenous abscisic acid alter hydrogen peroxide accumulation and differentially regulate the expression of two maize rd22-like genes. Scientific reports, 7: 1–12.

Prout T, Barker J. 1993. F statistics in drosophila buzzatii: selection, population size and inbreeding. Genetics, 134: 369–375.

R Core Team. 2020. R: A Language and Environment for Statistical Computing. R Foundation for Statistical Computing, Vienna, Austria.

Ravindran SP, Herrmann M, Cordellier M. 2019. Contrasting patterns of divergence at the regulatory and sequence level in european daphnia galeata natural populations. Ecology and evolution, 9: 2487–2504.

Roberge C, Guderley H, Bernatchez L. 2007. Genomewide identification of genes under directional selection: gene transcription qst scan in diverging atlantic salmon subpopulations. Genetics, 177: 1011–1022.

Rockman MV, Kruglyak L. 2006. Genetics of global gene expression. Nature Reviews Genetics, 7: 862.

Roelofs D, Overhein L, De Boer M, Janssens T, Van Straalen N. 2006. Additive genetic variation of transcriptional regulation: metallothionein expression in the soil insect orchesella cincta. Heredity, 96: 85.

Spitze K. 1993. Population structure in daphnia obtusa: quantitative genetic and allozymic variation. Genetics, 135: 367–374.

Swarts K, Gutaker RM, Benz B, Blake M, Bukowski R, Holland J, Kruse-Peeples M, Lepak N, Prim L, Romay MC et al. 2017. Genomic estimation of complex traits reveals ancient maize adaptation to temperate north america. Science, 357: 512–515.

Takeda S, Matsuoka M. 2008. Genetic approaches to crop improvement: responding to environmental and population changes. Nature Reviews Genetics, 9: 444.

Thompson R. 2008. Estimation of quantitative genetic parameters. Proceedings of the Royal Society B: Biological Sciences, 275: 679–686.

Walley JW, Sartor RC, Shen Z, Schmitz RJ, Wu KJ, Urich MA, Nery JR, Smith LG, Schnable JC, Ecker JR et al. 2016. Integration of omic networks in a developmental atlas of maize. Science, 353: 814–818.

Wang B, Lin Z, Li X, Zhao Y, Zhao B, Wu G, Ma X, Wang H, Xie Y, Li Q et al. 2020. Genome-wide selection and genetic improvement during modern maize breeding. Nature Genetics, pp. 1–7.

Wang RL, Stec A, Hey J, Lukens L, Doebley J. 1999. The limits of selection during maize domestication. Nature, 398: 236.

Whitehead A, Crawford DL. 2006. Neutral and adaptive variation in gene expression. Proceedings of the National Academy of Sciences, 103: 5425–5430.

Whitlock MC. 2008. Evolutionary inference from qst. Molecular ecology, 17: 1885–1896.

Wright SI, Bi IV, Schroeder SG, Yamasaki M, Doebley JF, McMullen MD, Gaut BS. 2005. The effects of artificial selection on the maize genome. Science, 308: 1310–1314.

Xu H, Li Y, Yan Y, Wang K, Gao Y, Hu Y. 2010. Genome-scale identification of soybean burp domain-containing genes and their expression under stress treatments. BMC plant biology, 10: 197.

Zhou P, Hirsch CN, Briggs SP, Springer NM. 2019. Dynamic patterns of gene expression additivity and regulatory variation throughout maize development. Molecular plant, 12: 410–425.

Zhou P, Li Z, Magnusson E, Cano FG, Crisp PA, Noshay JM, Grotewold E, Hirsch CN, Briggs SP, Springer NM. 2020. Meta gene regulatory networks in maize highlight functionally relevant regulatory interactions. The Plant Cell, 32: 1377–1396.

